# Application and Evaluation of Flipped Teaching Based on Video Conference in Standardized Training for Internal Medicine Residents

**DOI:** 10.1101/2022.04.17.488599

**Authors:** Xiao-Yu Zhang

## Abstract

**Background:** Infectious disease training was a necessary part of standardized training for internal medicine residents, and a designated hospital by the health administration department provided infectious diseases training for residents in those hospitals that did not meet the training standards for infectious diseases in the region. However, due to holidays, coordination of training dates among hospitals, and vacation of other reasons, the actual training time in the Department of Infectious Diseases might be insufficient.

**Objective:** I aimed to explore Flipped Teaching with Video Conference as the carrier in infectious disease training for internal medicine residents, to make up for the lack of actual training time of the Department of Infectious Diseases for those residents caused by vacation, and to ensure the smooth implementation and quality assurance of infectious disease training for those residents.

**Methods:** Vertical management mode was adopted, management team and lecturer team were established, and training program and teaching implementation were formulated. Flipped Teaching based on Video Conference was carried out for internal medicine residents of dispatched hospitals who planned to participate in infectious diseases training of the designated hospital in April. The quantitative analysis was applied to this teaching evaluation, and the evaluation indexes were included into statistical analysis to evaluate the effect of the teaching model.

**Results:** All 19-member internal medicine residents participated in the Flipped Teach based on Video Conference from April 1 to 4, of which 12 residents were scheduled to participate in infectious diseases training from March 1 to April 30, and 7 residents were scheduled to participate in infectious diseases training from April 1 to May 31. A management team of 6 internal medicine residents was built, and a group of 12 lecturers were composed of 12 those residents who were scheduled to receive infectious diseases training in the designated hospital from March 1 to April 30. According to the requirements of training diseases in the Department of Infectious Diseases, 12 training contents were selected to carry out teaching, and the implementation rate of teaching plan was over 90%. A total of 197 feedback questionnaires were collected. The feedback that the teaching quality was “good” and “very good” accounted for more than 96%, and the attendance rate of the whole teaching process reached more than 94%. About this teaching, 6 internal medicine residents put forward 18 suggestions of “Improvement suggestions”, accounting for 9.1%; and 11 internal medicine residents gave 110 suggestions of “Praise highlights”, accounting for 55.8%. The overall evaluation feedback of flipped teaching was good, P<0.001.

**Conclusion:** Flipped Teach based on Video Conference was generally effective in carrying out for internal medicine residents participating in the infectious diseases training, and it could be used as a supplementary training method for standardized training of internal medicine residents to make up for the shortage of actual training period in a certain stage.

## Introduction

In view of the rapid evolution of infectious diseases spectrum in Shanghai in recent years and the need for effective response to public health emergencies, the Department of Infectious Diseases was decided to be included in the compulsory rotation subject of standardized training for internal medicine residents, so as to further improve the knowledge structure of internal medicine residents and meet the social demand for the diagnosis and treatment of infectious diseases. For the training hospitals that did not have a ward of the Department of Infectious Diseases, or the training hospitals that could not meet the diseases required for the training of the Department of Infectious Diseases, internal medicine residents of which should be arranged to take part in the infectious diseases training of the designated hospital by the health administration department.[1]

With the continuous promotion of the infectious diseases training for internal medicine residents in Shanghai, more and more hospitals participated in the collaborative training of internal medicine residents in the designated hospital, and the number of internal medicine residents entering the infectious diseases training of the designated hospital gradually had been increasing. According to the requirements of the standardized training program for internal medicine residents, internal medicine residents attending the infectious diseases training in the designated hospital needed to carry out professional training for 1-2 months. In order to unify management and training, it was necessary to arrange a certain number of internal medicine residents to enter the designated hospital for infectious diseases training in an orderly manner every month, and make orderly connection so as to ensure the quality of training and rational use of public medical resources. During the operation of the project, the training cycle of internal medicine residents in the designated hospital for infectious diseases training might be insufficient, due to holidays or vacations, coordination of training dates between multiple hospitals, and other subjective and objective reasons. Therefore, I explored a new teaching mode to further improve the infectious diseases training for internal medicine residents.

In the Flip Teaching research report, the flipped classroom approach in health professions education yielded a significant improvement in student learning compared with traditional teaching methods[3], the flipped classroom and lecture were essentially equivalent[4], and an increased perceived value and acceptability of this model was noted by the participants[5]. Therefore, I explored the use of Video Conference as the carrier to carry out Flipped Teaching, and constructed the teaching mode of “Flipped Teaching in Standardized Training for Internal Medicine Residents Based on Video Conference” to perform distance online teaching. The mode was applied to the infectious diseases training for internal medicine residents, and preliminary evaluation was conducted.

## Objects and Methods

### Subjects

The study objects were internal medicine residents who planned to take training in the designated hospital in April. And they were also those residents who had participated in the standardized training for residents in Shanghai. Their dispatched hospital had signed an “Agreement on Joint Training of Internal Medicine Residents” with the designated hospital. The informed consents of participants in internal medicine residents were obtained, including their data being used for the training and the research, and that this study was conducted in accordance with the Declaration of Helsinki.

### Construction of Flipping Teaching Mode with Video Conference as the Carrier

The Flipped Teaching with Video Conference as the carrier adopted “vertical management mode” for management[2], and management team and lecturer team were established. According to the training requirements of the Department of Infectious Diseases in the Standardized Training Content and Standard for Resident Physicians (2021 Edition) -- Internal Medicine Training Rules, the management team would formulate the training program, and the lecturer team would select the training content and carry out teaching activities according to the plan. The following was the implementation of the teaching plan and the evaluation of the teaching work. The whole teaching organization, teaching implementation, discussion after teaching, teaching management and teaching evaluation were carried out online without restriction of physical space.

### Evaluation Indexes and Criteria

This teaching model was evaluated from four aspects, including the implementation of the teaching plan, the attendance of the Flipped Teaching, the evaluation of teaching quality, and the overall evaluation of teaching.

Six teaching plan indicators (teaching on the planned time, teaching on the planned content, making PPT fully, providing references, unifying the teaching content and training program, and participating in after-class discussion) were established to evaluate the implementation of the teaching plan. Three attendance indicators (online on time, middle roll call and end on time) were established to evaluate the online attendance of the Flipped Teaching were completed. Nine teaching quality indicators (rigorous teaching attitude, punctual class, detailed and accurate teaching content, reasonable structure and clear process, highlighting teaching key points, clear teaching difficulties, accurate and refined language, combining theory with clinical practice, improving ability to analyze and deal with the disease) were established to evaluate the teaching quality. The overall evaluation of teaching adopted open questionnaire to evaluate each teaching without limit.

The teaching plan indicators and the attendance indicators were completed by the organizer, and the teaching quality indicators and the overall evaluation content were completed by every trainee for each teaching session. Among them, teaching plan indicators and teaching quality indicators were objective indicators, and they were derived from Teaching Evaluation Table of Standardized Residency Training; while overall evaluation indicators was subjective. The feedback of the over evaluation from open questionnaire was firstly classified according to the evaluation content. The details of classification as follow: If the content of a questionnaire feedback was pointing out deficiencies or needing improvement of one teaching session, this feedback was classified as “Improvement suggestions”; If all the content of a questionnaire feedback was praising highlights or learning achievements of one teaching session, this feedback was classified as “Praise highlights”; If the content of a questionnaire feedback was no special suggestions of one teaching session, this feedback was classified as “No special suggestions”.

### Software Application and Statistical Analysis

Video Conference adopted Tencent Conference software to carry out the Flipped Teaching, including: teaching organization, teaching implementation, discussion after teaching, teaching management, and teaching evaluation.

The Questionnaire Star software was used to develop teaching quality indicators and overall evaluation indicators, and carry out star survey after class; and then the data of above indexes were downloaded from the software and incorporated into statistical analysis.

SPSS software version 23.0 (SPSS Inc. Chicago, IL, USA) was used for statistical analysis of the data. The data conforming to normal distribution were expressed as mean ± standard deviation to reflect the distribution of the study indicators. The counting data was represented by example (%) to reflect the composition ratio of the study indicators. Pearson chi-square test was used for the counting data. A P value of two-sided less than 0.05 was considered as statistically significant.

## Results

### Basic Information of Physicians Participating in Teaching Activities

A total of 19 internal medicine residents participated in the Flipped Teaching program, all from tertiary hospitals. Among them, 9 were male, accounting for 47.4%. The average age was 29.5 years. 5 had bachelor degree, accounting for 26.3%; 4 had master degree, accounting for 21.1%; 10 had doctor degree, accounting for 52.6%. 17 internal medicine residents were qualified as practicing physicians, accounting for 89.5%. 2 in the first year of training, accounting for 10.5%. 12 in the second year of training, accounting for 63.2%. 5 in the third year of training, accounting for 26.3%. 12 were scheduled to participate in infectious diseases training in the designated hospital from March 1 to April 30, accounting for 63.1%; 7 were scheduled to participate in infectious diseases training in the designated hospital from April 1 to May 31, accounting for 36.8%. The detailed information is shown in **Table 1**.

**Table 1.**
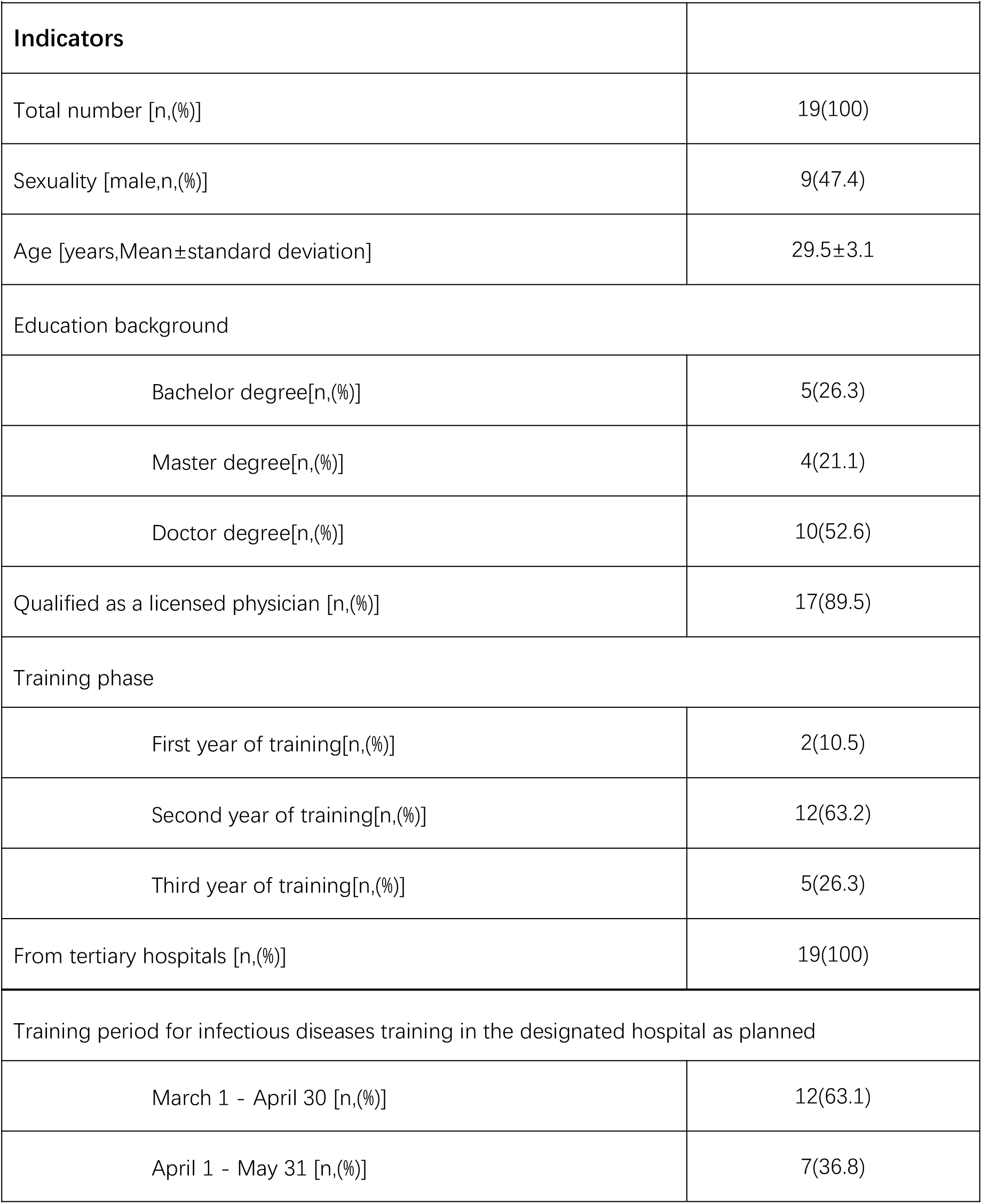
Baseline of Internal Medicine Residents Participating in Flipped Teaching

|  |  |
| --- | --- |
| Indicators |  |
| Total number [n,(%)] | 19(100) |
| Sexuality [male,n,(%)] | 9(47.4) |
| Age [years,Mean±standard deviation] | 29.5±3.1 |
| Education background |  |
| Bachelor degree[n,(%)] | 5(26.3) |
| Master degree[n,(%)] | 4(21.1) |
| Doctor degree[n,(%)] | 10(52.6) |
| Qualified as a licensed physician [n,(%)] | 17(89.5) |
| Training phase |  |
| First year of training[n,(%)] | 2(10.5) |
| Second year of training[n,(%)] | 12(63.2) |
| Third year of training[n,(%)] | 5(26.3) |
| From tertiary hospitals [n,(%)] | 19(100) |
| Training period for infectious diseases training in the designated hospital as planned |  |
| March 1 - April 30 [n,(%)] | 12(63.1) |
| April 1 - May 31 [n,(%)] | 7(36.8) |

### Flipped Teaching Organization

The Flipped Teaching with Video Conference as the carrier was organized and implemented by the contact person of the “Rotation Training of Infectious Diseases Department for Internal Medicine Residents” project, which was managed by the “vertical management mode”. The management team was formed together with the monitor and group leader of this training course, and 12 lecturers were composed of those residents who were scheduled to take part in the infectious diseases training of the designated hospital from March 1 to April 30.

The contents of the lecture were as follows: Analysis of Chronic Hepatitis B, Study on Guidelines for Prevention and Treatment of Hepatitis C (2019 Edition), Identification and Treatment of Clostridium Difficile Associated Diarrhea, Bacterial Liver Abscess, Diagnosis and Treatment of Tuberculosis, Guidelines for the Diagnosis and Treatment of Syphilis, Diagnosis and Treatment of Tuberculous Meningitis, Infective Endocarditis, Cryptococcal Meningitis, Study on AIDS Diagnosis and Treatment Guide in China (2021 edition), Diagnosis and Treatment of Cirrhotic Ascites, and Diagnosis and Treatment of liver Failure. Teaching tasks were assigned to those residents according to the time period, and PPT was developed to carry out teaching activities according to clinical guidelines. The contents of the lecture are shown in **Table 2**.

**Table 2.** Flipped Teaching Plan

| **Table 2** Flipped Teaching Plan |  |  |
| --- | --- | --- |
| **Teaching date** | **Teaching time** | **content of courses** |
| April 1 | 13:35-14:20 | Analysis of Chronic Hepatitis B |
| April 1 | 14:35-15:20 | Study on Guidelines for Prevention and Treatment of Hepatitis C (2019 Edition) |
| April 1 | 15:35-16:20 | Identification and Treatment of Clostridium Difficile Associated Diarrhea |
| April 2 | 13:35-14:20 | Bacterial Liver Abscess |
| April 2 | 14:35-15:20 | Diagnosis and Treatment of Tuberculosis |
| April 2 | 15:35-16:20 | Guidelines for the Diagnosis and Treatment of Syphilis |
| April 3 | 13:35-14:20 | Diagnosis and Treatment of Tuberculous Meningitis |
| April 3 | 14:35-15:20 | Infective Endocarditis |
| April 3 | 15:35-16:20 | Cryptococcal Meningitis |
| April 4 | 13:35-14:20 | Study on AIDS Diagnosis and Treatment Guide in China (2021 edition) |
| April 4 | 14:35-15:20 | Diagnosis and Treatment of Cirrhotic Ascites |
| April 4 | 15:35-16:20 | Diagnosis and Treatment of liver Failure |

### Evaluation of Teaching Plan Implementation

The Flipped Teaching was carried out according to plan, with real-time online management and evaluation. In the teaching process, one of the lecturers delayed the start time of teaching on the planned time, because he was not familiar with Video Conferencing software; and the rest of the lecturers carried out teaching activities on time. Teaching plan met the requirements for 11 times, accounting for 91.7%. All the other teaching plan indicators were in compliance with the compliance rate of 100%. The detailed information is shown in **Table 3**.

**Table 3.** Evaluation of Teaching Plan Implementation

| **Table 3** Evaluation of Teaching Plan Implementation |  |  |  |
| --- | --- | --- | --- |
| **Teaching plan indicators** | **CT** | **NCT** | **CR (%)** |
| Teaching on the planned time | 11 | 1 | 91.7 |
| Teaching on the planned content | 12 | 0 | 100 |
| Making PPT fully | 12 | 0 | 100 |
| Providing references | 12 | 0 | 100 |
| Unifying the teaching content and training program | 12 | 0 | 100 |
| Participating in after-class discussion | 12 | 0 | 100 |

**Abbreviations:** CT, Coincidence times ; NCT, Not-Coincidence times; CR, Coincidence rate.
**Note:** Coincidence rate (%) = Coincidence times of teaching indicators / Total times of teaching indicators \*100%.

### Teaching Attendance and Feedback

The whole process of attendance was checked for this Flipped Teaching based on Video Conference, and the three time nodes of “online on time”, “middle roll call” and “end on time” were included in the statistics. There were 18, 18 and 19 internal medicine residents in attendance at the above three time nodes, and the attendance rates were 94.7%, 94.7% and 100% respectively. One of those residents failed to go online on time because he was not familiar with Video Conference software, and one asked for leave and went offline due to an emergency. The detailed attendance is shown in **Table 4**.

**Table 4.** Attendance of Flipped Teaching

| **Table 4** Attendance of Flipped Teaching |  |  |
| --- | --- | --- |
| **Attendance indicators** | **Residents in attendance** | **Attendance rates(%)** |
| Online on time | 18 | 94.7 |
| Middle roll call | 18 | 94.7 |
| End on time | 19 | 100 |

**Note:** Attendance rate (%) = Number of internal medicine residents in attendance / Total number of internal medicine residents of this Flipped Teaching for the infectious training \*100%

### Quality Evaluation of Teaching

The teaching quality of the Flipped Teaching based on Video Conference was investigated from the questionnaire star, and 179 effective feedback questionnaires were collected. Feedback “good” of nine indicators (“Rigorous teaching attitude”, “Punctual class”, “Detailed and accurate teaching content”, “Reasonable structure and clear process”, “Highlighting teaching key points”, “Clear teaching difficulties”, “Accurate and refined language”, “Combining theory with clinical practice”, and “Improving ability to analyze and deal with the disease”) was 26, 29, 36, 33, 32, 33, 31, 28 and 30, accounting for 13.2%, 14.7%, 18.3%, 16.8%, 16.3%, 16.8%, 15.8%, 14.3% and 15.3%, respectively. And Feedback “very good” of the above nine indicators 170, 167, 157, 159, 162, 157, 161, 163 and 163, accounting for 86.3%, 84.8%, 79.7%, 80.7%, 82.2%, 79.7%, 81.7%, 82.7% and 82.7%, respectively. Overall, teaching quality was rated as “good” and “very good” by more than 96%. The detailed evaluation of teaching quality is shown in **Table 5**.

**Table 5.** Quality Evaluation of Flipped Teaching

| **Table 5** Quality Evaluation of Flipped Teaching |  |  |  |  |  |
| --- | --- | --- | --- | --- | --- |
| Teaching quality indicators[n,(%)] | VP(%) | PR(%) | GN(%) | GD(%) | VG(%) |
| Rigorous teaching attitude | 0(0) | 0(0) | 1(0.5) | 26(13.2) | 170(86.3) |
| Punctual class | 0(0) | 0(0) | 1(0.5) | 29(14.7) | 167(84.8) |
| Detailed and accurate teaching content | 0(0) | 0(0) | 4(2.0) | 36(18.3) | 157(79.7) |
| Reasonable structure and clear process | 0(0) | 0(0) | 5(2.5) | 33(16.8) | 159(80.7) |
| Highlighting teaching key points | 0(0) | 0(0) | 3(1.5) | 32(16.3) | 162(82.2) |
| Clear teaching difficulties | 0(0) | 0(0) | 7(3.5) | 33(16.8) | 157(79.7) |
| Accurate and refined language | 0(0) | 0(0) | 5(2.5) | 31(15.8) | 161(81.7) |
| Combining theory with clinical practice | 0(0) | 0(0) | 6(3.0) | 28(14.3) | 163(82.7) |
| Improving ability to analyze and deal with the disease | 0(0) | 0(0) | 4(2.0) | 30(15.3) | 163(82.7) |

**Abbreviations:** VP, Very poor; PR, Poor; GR, General; GD, Good; VG, Very good.
**Note :** 197 feedback of teaching quality evaluation were collected through questionnaires.

### Overall Evaluation of Teaching

In the overall evaluation of the Flipped Teaching based on Video Conference, 6 internal medicine residents filled in “Improvement suggestions” feedback in 18 questionnaires, accounting for 9.1%; 11 internal medicine residents proposed “Praise highlights” feedback in 110 questionnaires, accounting for 55.8%. 10 internal medicine residents had “No special suggestions” feedback in 69 questionnaires, accounting for 35.0%. The overall evaluation feedback of Flipped Teaching based on Video Conference was good, P<0.001. The detailed overall evaluation of feedback is shown in **Table 6**.

**Table 6.**
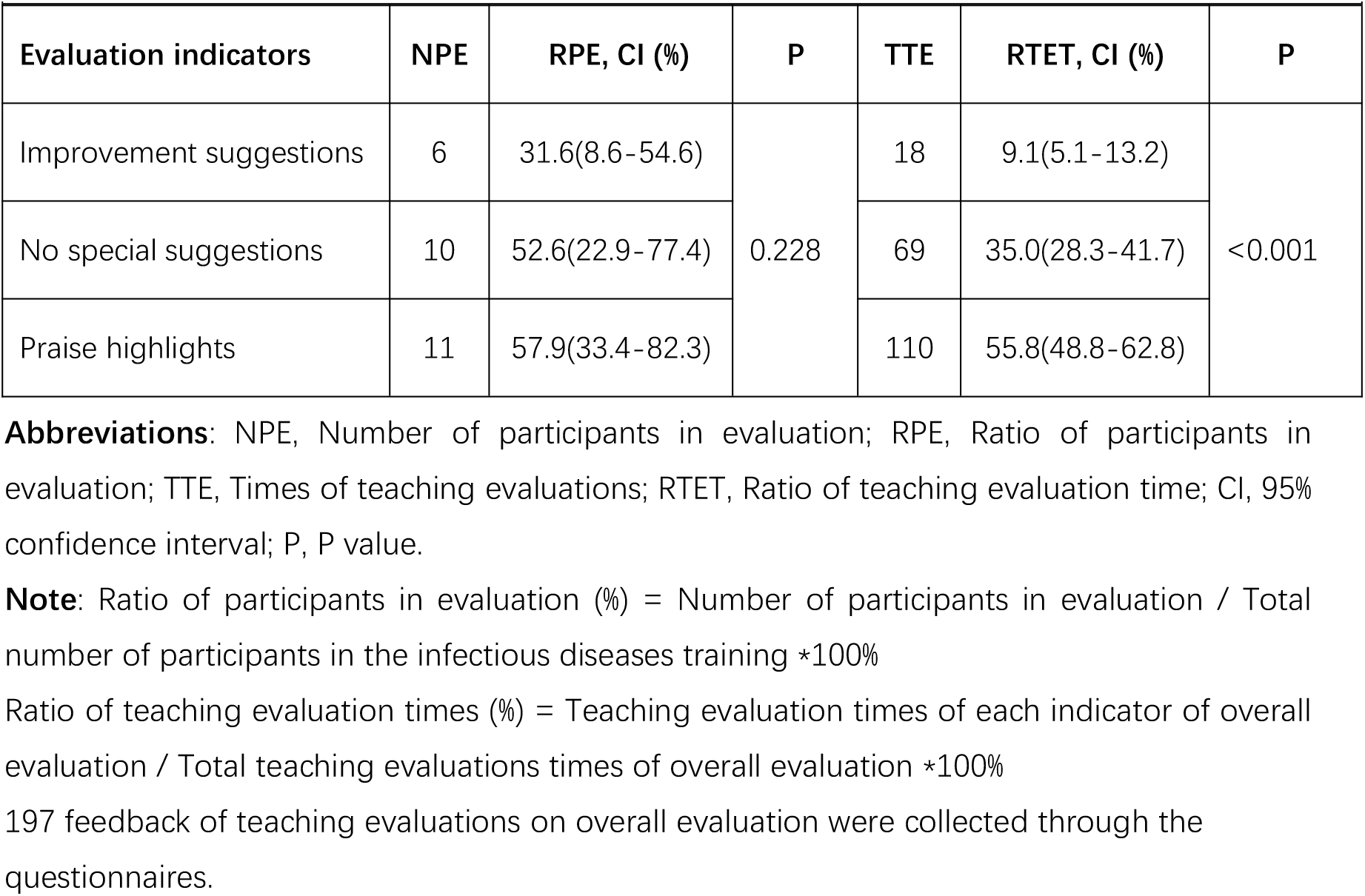
Overall Evaluation of Flipped Teaching

## Discussion

The participation of internal medicine residents participating in the infectious diseases training was carried out under specific conditions, which was the perfection of the knowledge structure of the standardized training of internal medicine residents and the social needs for the diagnosis and treatment of infectious diseases. Those training hospitals that did not meet the training standards for infectious diseases should cooperate with the designated hospital to conduct infectious diseases training for internal medicine residents. Due to the fact that the internal medicine residents who participated in the infectious diseases training of the designated came from more than 10 hospitals in Shanghai, the personnel management, and implementation plan and training progress of internal medicine residents in each hospital were different to some extent. In order to ensure the quality of internal medicine residents training and rationally allocate the training resources, a certain number of internal medicine residents were accepted to participate in the infectious diseases training of the designated hospital every month. In the actual operation, due to involve legal holidays, vocations for other reasons and emergencies; some of the internal medicine residents taking part in the infectious diseases training of the designated hospital could not meet the training time requirements. To this end, I applied Flipped Teaching based on Video Conference to carry out infectious diseases training for internal medicine residents. Therefore, it was obviously advanced and necessary to carry out the application and research of this teaching model.

Flipped classrooms showed many advantages when tested in a radiology classroom setting, making up for some inadequacies of didactic classrooms; but it was needed to make improvements to make it more suitable for the Chinese medical education mode.[6] The flipped classroom approach showed promise in ophthalmology clerkship teaching; but, it had some drawbacks; further evaluation and modifications were required before it could be widely accepted and implemented.[7] Based on the above, I carried out this teaching model research. The whole Flipped Teaching mode involved organizational structure, training content, training plan, training implementation, training management and evaluation. This teaching research was a prospective study, establishing organizational structure and management model, and developing training content and training plan. The implementation and quality analysis of the teaching model was evaluated by assessment and after-school questionnaire survey. Then the practicability, rationality, feasibility and effectiveness of this teaching model were evaluated. Therefore, this teaching research had distinct scientific nature.

The teaching organization structure included training organization, management mode, management team and lecturer team. This training was organized and implemented by the contact person of the “Rotation Training of Infectious Diseases Department for Internal Medicine Residents” project, to ensure the cooperation and support of teaching administrative departments and internal medicine residents of each hospital dispatched, and to mobilize the enthusiasm of internal medicine residents for this infectious diseases training project. A management team of six was set up in this teaching work, which were used to assist the project contact person to organize and implement this teaching activity, and supervised the teaching plan and supporting work, In order to further develop the those residents’ subjective initiative. A total of 12 lecturers were organized, consisting of those residents who planned to attend training on the infectious diseases in the designated hospital from March 1 to April 30; As a result, they had one month of clinical training experience in the Department of Infectious Diseases. They chose target diseases to give lectures based on their own clinical practice and shared their learning experience of this disease. In this way, the teaching could encourage the lecturers to learn and summarize the clinical knowledge of the selected diseases. 12 lecturers were selected in this teaching to encourage more qualified lecturers to participate in teaching activities and ensure the coverage of the teaching content. This teaching adopted vertical management mode. Since the “vertical management” mode had gained good experience and evaluation in the field of standardized training for public health physicians[2], this mode was adopted in this teaching organization, which also reflected that the teaching organization had very good practicability.

The content of this teaching was based on the establishment of the training program of training syllabus for internal medicine residents, with the orientation of the essential diseases and key diseases in the infectious diseases training. The goal of this teaching was to train the diagnosis and treatment skills of infectious diseases for internal medicine residents. The main contents were based on the clinical diagnosis and treatment guidelines of infectious diseases. All above was to ensure that the teaching purpose was consistent with requirements of the infectious diseases training, and the standardized training of internal medicine residents. So, 12 diseases were selected as the main topics of the teaching, covering all the diseases that must be mastered, and involving other key diseases of the infectious diseases training. Therefore, the teaching and training content was reasonably designed, which met the needs of the infectious diseases training at the level of internal medicine residents and reflected the rationality of the teaching mode.

In the part of planning implementation, it was an important part of teaching work to formulate training plans and organize implementation under the condition of definite training content. In order to carry out the teaching contents in an orderly manner, targeted teaching plans were drawn up, and the teaching contents needed to be implemented for specific personnel and specific time periods, so that the lecturers and participating residents could make full preparations. The management group made a good teaching time frame; while and the lecturers took the initiative to participate, and selected their own lecturing content and lecturing time. Once the content of the lecture was determined, no modification would be made unless in an emergency situation. If the teaching could not be carried out within the planned time, the lecturer’s teaching should be adjusted to the last time period in an orderly manner. The original plan of the 12 disease teaching tasks was to be completed in 4 days, and three time periods were arranged for orderly teaching activities every afternoon. In the implementation of the teaching plan, it was adopted for unified planning, group management, classified hosting, punctual teaching, online discussion, whole attendance, and “Questionnaire Star” survey feedback after class. In this teaching activity, one of the lecturers delayed the start time of teaching, because he was not familiar with Video Conference software. Therefore, in the formal organization of teaching activities, teaching preparation should be done more carefully for every section. For example, software drills and trial lectures should be carried out in the early stage of teaching. In the process of teaching implementation, the proportion of “Teaching on the planned time” was 91.7%, and the proportion of “Teaching on the planned content”, “PPT making fully”, “Providing references”, “Unifying teaching content and training outline” and “Participation in after-class questions” was 100%. From the perspective of teaching arrangement and implementation, this teaching plan had been implemented smoothly and has strong feasibility.

This Flipped Teaching based on Video Conference adopted online whole-process monitoring, and three time nodes of “On-time online”, “Middle roll Call” and “On-time end” were included in the statistics. The attendance rate was 94.7%, 94.7% and 100% respectively. One of them failed to go online on time because he was not familiar with the Video Conference software, and one of them asked for leave and went offline because of emergency during the process. From the statistics of the above three time nodes, whole-process monitoring was helpful to stabilize the teaching attendance rate. It was also conducive to mastering the teaching of emergency, timely discovery, timely treatment. From the perspective of teaching management experience, online questioning could motivate the lecturers to prepare for teaching, promote the lecturers to take the initiative to learn for knowledge reserve, and mobilize the active learning consciousness and enthusiasm of those residents. And the attendance of this teaching model was higher than that of other types of flipped classes, where attendance was 30-80%[8]. Through the Questionnaire Star to the teaching quality of 9 indicators feedback, the results showed that the teaching quality was “Good” and “Very good” accounting for more than 96%. In other specialized clinical training, flipped classroom was also well received and preferred, and it improved teaching satisfaction.[9–11] In terms of the attendance and teaching quality evaluation of internal medicine residents, the effectiveness of this teaching model was relatively good.

In the overall evaluation of this teaching, 11 internal medicine residents raised “Praise highlights” in 110 questionnaires, accounting for 55.8%; which was higher than other feedback, P<0.05. It reflected that the Flipped Teaching based on Video Conference was generally recognized, the opinion which was consistent with other clinical training studies that a flipped classroom approach in physiotherapy education resulted in improved student performances in this professional programme[12], even a study suggested flipped classroom for cardiovascular prevention curriculum showed greater effectiveness in knowledge gain[13]. However, 10 internal medicine residents had “No special suggestions” feedback among the 69 questionnaires; and 6 internal medicine residents filled in “Improvement suggestions” feedback in 18 questionnaires, accounting for 9.1%. The above feedback also suggested that the teaching mode still needed to be improved. In promoting the application of this mode, the organizer also needed to follow up the training work in real time, collected the feedback of those residents, and made continuous improvement based on the requirements of infectious diseases knowledge in the standardized training for internal medicine residents.

## Conclusions and Perspectives

The Flipped Teaching with Video Conference as carrier for internal medicine residents participating in the infectious diseases training was generally effective. The degree of participation and recognition of those residents in this Flipped Teaching was relatively good, and the implementation of teaching program was feasible. Application of this teaching mode could make up for the shortage of actual training time of residents in a certain stage.

## Ethical Approval and Consent to Participate

Informed consents of participants in the standardized training for internal medicine residents were obtained for the training and the study. The study received Institutional Review Board (IRB) approval by the Shanghai Public Health Clinical Center Ethics Committee. The IRB number was No. 2021-S026-01.

## Acknowledgments

This study was supported by the internal medicine residents who had participated in the standardized training for residents in Shanghai. Thanks to the teaching administration department and the resident standardized training base of the united training hospital for their support. Thanks for the support of Shanghai and National administration departments of standardized training for resident physicians.

## Authors’ Contributions

Xiao-Yu Zhang made conception, design, acquisition of data, analysis and interpretation of data, drafted and revised the manuscript, and agreed to be accountable for all aspects of the work.

## Consent for Publication

The author has read and agreed to the published version of the manuscript.

## Data Availability Statement

The data included in the manuscript submitted to the journal is transparent,and all relevant data is available within the tables in the publication.

